# Influenza infected macrophages release viral ribonucleoproteins that shape the local host response

**DOI:** 10.1101/2023.12.04.570004

**Authors:** Svenja Fritzlar, Seema S. Lakdawala, Jason A. Roberts, Lachlan J. M. Coin, Yi-Mo Deng, Jean Moselen, Valerie M. Le Sage, Jason M. Mackenzie, Larisa I. Labzin, Andrew G. Brooks, Patrick C. Reading, Sarah L. Londrigan

## Abstract

Airway epithelial cells and macrophages (MΦ) represent cellular targets of infection by influenza A virus (IAV). Epithelial cells support IAV infection and replication by enabling the generation and release of new viral particles (productive replication). In contrast, MΦ are susceptible to initial IAV infection but the release of infectious viral particles is inhibited through abortive replication. Despite the lack of infectious virions released from infected MΦ, we detected newly synthesised viral RNA and nucleoprotein (NP) in MΦ supernatants. We show that viral RNA is released from infected MΦ as viral ribonucleoprotein (vRNP) complexes which elicits potent inflammatory responses when exposed to uninfected cells. These vRNPs specifically induced IL-1β, CXCL13, IL-32, CCL4 and CXCL10 in uninfected cells and were at least partially sensed through the RIG-I/MDA5 pathway. While MΦ represent a dead-end for IAV infection through abortive replication, the release of vRNPs shapes immune repsonses of uninfected cells in the local microenvironment.

## Introduction

Airway macrophages (MΦ) play a crucial role in the host defence against respiratory virus infections and are abundant in the respiratory tract. MΦ phagocytose virus-infected cells, which has been shown to significantly decrease the viral burden during infection and help in viral clearance ^1–3^. MΦ can also be directly infected with viruses, where viral pathogen-associated molecular patterns (PAMPs) including viral RNA are sensed through retinoic acid-inducible gene I (RIG-I) and toll like receptors (TLRs) ^4–8^. This influences MΦ-mediated immune signalling and antiviral responses resulting in the release of pro-inflammatory cytokines and type I interferons (IFNs) ^3,8^. The importance of protective MΦ responses in respiratory virus infection has also been demonstrated through chlodronate depletion of MΦ in animal models. MΦ-depleted mice infected with influenza A virus (IAV) and respiratory syncytial virus (RSV) displayed increased viral replication, enhanced pulmonary pathology and increased morbidity ^9–11^. MΦ-depleted ferrets and pigs infected with IAV also displayed increased disease severity ^12,13^. During IAV infection MΦ responses are predominantly protective and rely on tightly regulated antiviral and pro-inflammatory signatures to shape the early responses to infection ^14–16^. An increased understanding of the host responses that control IAV infection could provide opportunity for the development of novel host-directed antiviral strategies to combat severe IAV-induced disease.

Both airway epithelial cells and MΦ are susceptible to initial IAV infection, but only epithelial cells support productive replication defined by the formation and release of newly synthesised virions. This includes the synthesis of eight viral RNA (vRNA) segments (PA, PB1, PB2, HA, NP, NA, M and NS) packaged into viral ribonucleoprotein (vRNP). The vRNP comprises the vRNA in association with the viral nucleoprotein (NP) and polymerase proteins PA, PB1 and PB2. In contrast, mouse MΦ are susceptible to the early stages of infection with seasonal IAV but this results in an abortive replication without the release of infectious viral particles ^17,18^. We previously showed during abortive IAV replication in mouse MΦ that vRNA and mRNA are transcribed for all eight RNA segments and viral proteins are translated, but the assembly of infectious viral particles is blocked during the late stages of replication ^18^. Furthermore, type I IFN responses are critical for limiting IAV replication, because IAV infection of IFN-α/β receptor deficient mouse MΦ resulted in productive replication ^17^. In contrast, some highly pathogenic avian influenza viruses (HPAI) can overcome this block and productively replicate in MΦ ^19,20^, which has been mapped to the IAV hemagglutinin (HA) protein ^21^. These studies, along with *in vivo* animal studies where MΦ were depleted ^11^, suggest that MΦ present a dead end for seasonal IAV replication, limiting viral spread and reducing viral induced pathology within the airways.

Herein, we infected mouse MΦ with seasonal IAV and showed that during abortive replication, vRNPs are assembled and released contributing to significant levels of vRNA and NP in the supernatants of infected MΦ. These vRNPs were immunoreactive inducing potent inflammatory responses when exposed to uninfected cells. These findings reveal novel mechanistic insights into how MΦ shape the local host responses during IAV infection.

## Results

### Viral RNA and protein are released from IAV-infected macrophages during abortive replication

We infected mouse MΦ, including primary MΦ isolated from peritoneal exudate or the RAW264.7 cell line, with seasonal IAV to further investigate the block in late-stage replication. As expected, there was no significant increase in infectious viral titres in cell-free supernatants (SN) harvested at 2 vs. 24 hours post infection (hpi), which is consistent with abortive replication (Fig. 1A (i/ii)). In contrast, epithelial cells (Madin Darby Canine Kidney (MDCK) or murine lung (LA-4) cells) supported productive replication evidenced by a significant increase in viral titres in the SN between 2 and 24 hpi (Fig. 1A (i/ii)). Confocal microscopy analysis of newly synthesised IAV proteins produced during abortive replication in RAW264.7 MΦ confirmed susceptibility to the early stages of infection where the viral NP, polymerase (PA and PB2) and non-structural 1 and 2 (NS1 and NS2) proteins localised to cellular compartments similar to those in productively infected epithelial cells (Fig. SI 1A). Additionally, IAV proteins involved in late stages of infection such as HA and M2 trafficked to the cell surface where they co-localised (Fig. SI 1B). These observations are consistent with a defect in the late stages of IAV replication in MΦ involving viral assembly and release. Despite the absence of infectious viral particles in SN harvested from infected MΦ, a significant increase in vRNA, represented by the matrix (M) gene, was detected between 2 and 24 hpi in both RAW264.7 MΦ (Fig. 1B (i)) and primary MΦ SN (Fig. 1B (ii)). SN harvested from productively infected epithelial cells also showed a significant increase in M gene and other vRNA at 24 hpi, consistent with release of newly synthesised viral particles (Fig. 1B (i/ii), Fig.SI. 2A-C)). The release of vRNAs (represented by M gene) was not significantly reduced by treatment of cells with inhibitors blocking apoptosis and necroptosis (Fig. SI 2D) indicating that vRNA release is independent of these pathways. To assess if the presence of vRNA in MΦ SN was associated with significant perturbations in the abundance of each viral gene segment during abortive infection, we performed next-generation sequencing (NGS) on viral RNA isolated from infected RAW264.7 MΦ and epithelial cell (MDCK) SN (Fig. 1C). We confirmed the presence of all eight IAV RNA segments in SN obtained from infected MΦ and epithelial cells, where only minor differences in the abundance of specific IAV RNA segments were identified between the different cell types. We then extended our analysis to assess the presence of specific viral proteins. As expected, increasing levels of NP were detected in SN of IAV-infected epithelial cells over time, consistent with the release of viral particles during productive replication (Fig. 1D). Surprisingly, we also detected significant amounts of viral NP in the SN of IAV infected MΦ (RAW264.7 and primary MΦ) during abortive replication. The levels of NP detected in SN of infected MΦ were consistently higher than those in the SN of productively infected IAV-infected epithelial cells (Fig. 1D). Low levels of IAV HA and NA proteins were also detected in the SN of infected MΦ (data not shown). This indicates that substantial amounts of viral protein and viral RNA are released from IAV infected MΦ during abortive replication that is not associated with infectious viral particles.

**Figure 1:**
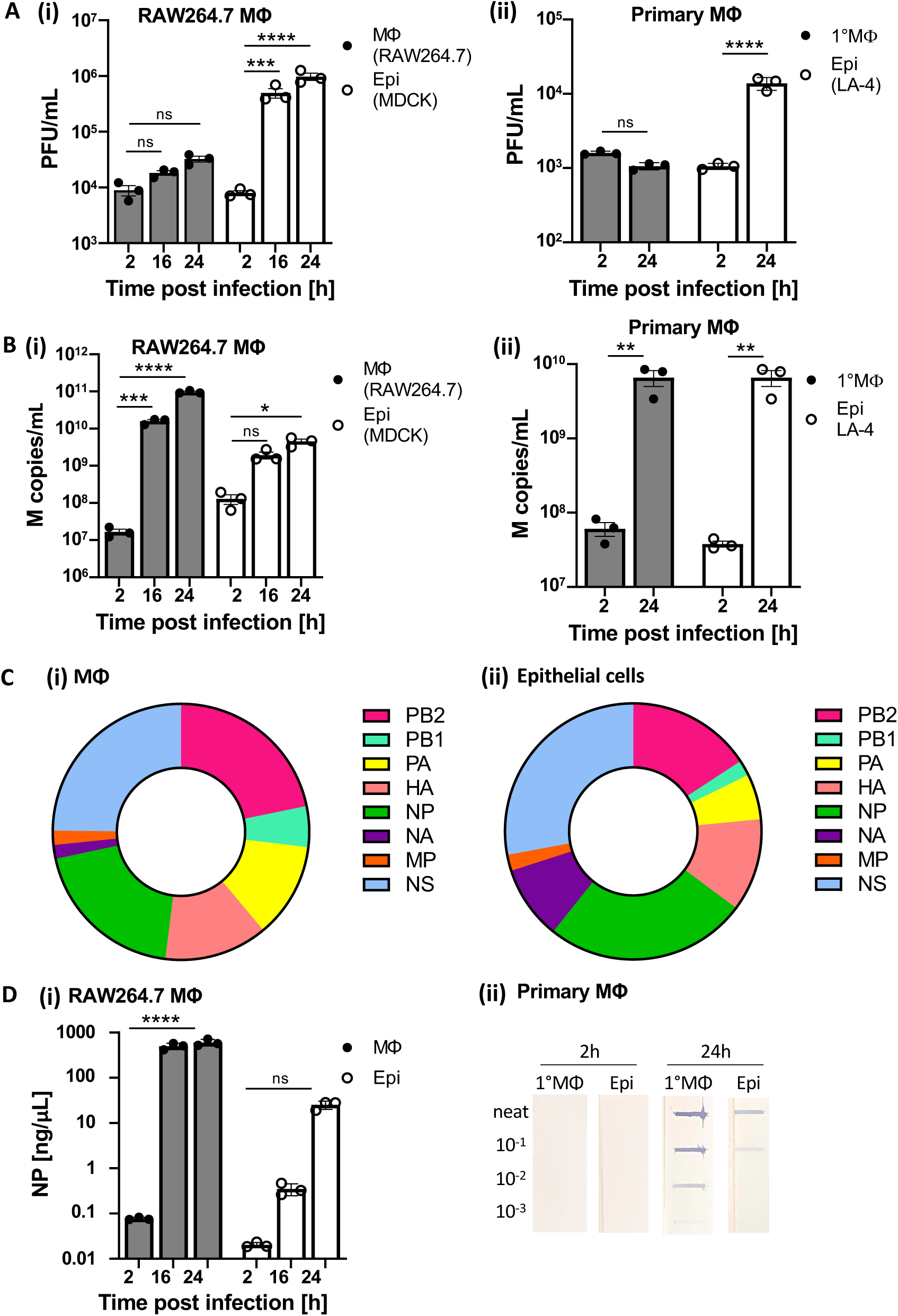
Viral RNA and protein are released from IAV-infected macrophages during abortive replication. Mouse MΦ (RAW264.7 or primary mouse MΦ, grey bars) and control epithelial cells (MDCK or LA-4 cells, white bars) were infected with IAV (MOI 5) and cell-free culture SN was analysed at 2, 16 and 24 hpi (RAW264.7 vs MDCK) or 2 and 24 hpi (primary mouse MΦ vs LA-4 cells). Release of infectious viral particles was assessed via **(A) (i/ii)** plaque assay (PFU/mL) and vRNA copies in the cell-free SN were analysed by **(B) (i/ii)** RT-qPCR (M copies/mL). **(C)** NGS was performed on cell-free SN of infected MΦ (RAW264.7) and epithelial cells (MDCK) 24 hpi. Graphs represent the average percentage (n=3) of each RNA segment out of total amount of RNA detected. The sequences for all eight influenza A genes were identical to the parent virus sequence. The presence of IAV NP in the cell-free SN was determined via **(D) (i)** IAV NP-specific AlphaLISA assay (ng/mL) or **(D) (ii)** slot blot (neat, 10^-1^, 10^-2^, 10^-3^ dilutions) analysis at 2 and 24 hpi. n=3 for all experiments. ANOVA or t-test used with p<0.5 *, p<0.1 **, p<0.01 ***, p<0.001 ****, ns: not significant.

### vRNA released from IAV infected macrophages is packaged into vRNP complexes

Next, we examined SN from IAV-infected MΦ and epithelial cells by transmission electron microscopy (TEM) (Fig. 2A (i/ii) and Fig. 2B). As expected, virions were easily and frequently observed in SN from productively infected epithelial cells at 24 hpi (Fig. 2B). While virions were absent from MΦ SN, we were intrigued to observe abundant helical structures indicative of vRNA complexed with viral proteins as vRNPs ^22^ (Fig. 2B). To further investigate the nature of the vRNP-like structures, we infected MΦ and epithelial cells with IAV for 24h and depleted residual input viral particles, as well as newly synthesised viral particles released during productive replication in epithelial cells, using red blood cells (RBC). SN samples were then immunogold-labelled using antibodies specific for IAV NP and analysed via TEM (Fig. 2A(iv)). IAV-infected MΦ SN contained RNA-like structures which clustered with IAV NP-labelled gold particles (Fig. 2C). We observed aggregate structures in virus-depleted epithelial cell SN, but these did not exhibit the typical helical structure of vRNPs. Although the epithelial cell SN did contain some gold particles, these were sporadic and did not specifically bind to the aggregate structures suggesting that vRNP complexes, outside of viral particles, were not present. Next, we used a co-precipitation approach to confirm both the presence and composition of the vRNP complexes in SN from IAV-infected MΦ but not epithelial cells. We immunoprecipitated the IAV PA protein and looked for association with either PB2 or NP as evidence of vRNP complexes by western blotting (Fig. 2A (v)). SN from IAV infected MΦ, but not epithelial cells showed co-precipitation of IAV PA with both the PB2 and NP proteins (Fig. 2D), indicating the presence of vRNP complexes in the SN of IAV infected MΦ during abortive replication. vRNPs were absent in the SN of productively infected epithelial cell controls, because they are packaged into viral particles and are therefore absent in the virus-depleted SN of IAV infected epithelial cells. Other IAV proteins that are not part of the vRNP complex, including the non-structural proteins NS1 and NS2, did not co-precipitate with the IAV PA protein. We conclude that IAV vRNP complexes are released from MΦ during abortive IAV replication.

**Figure 2:**
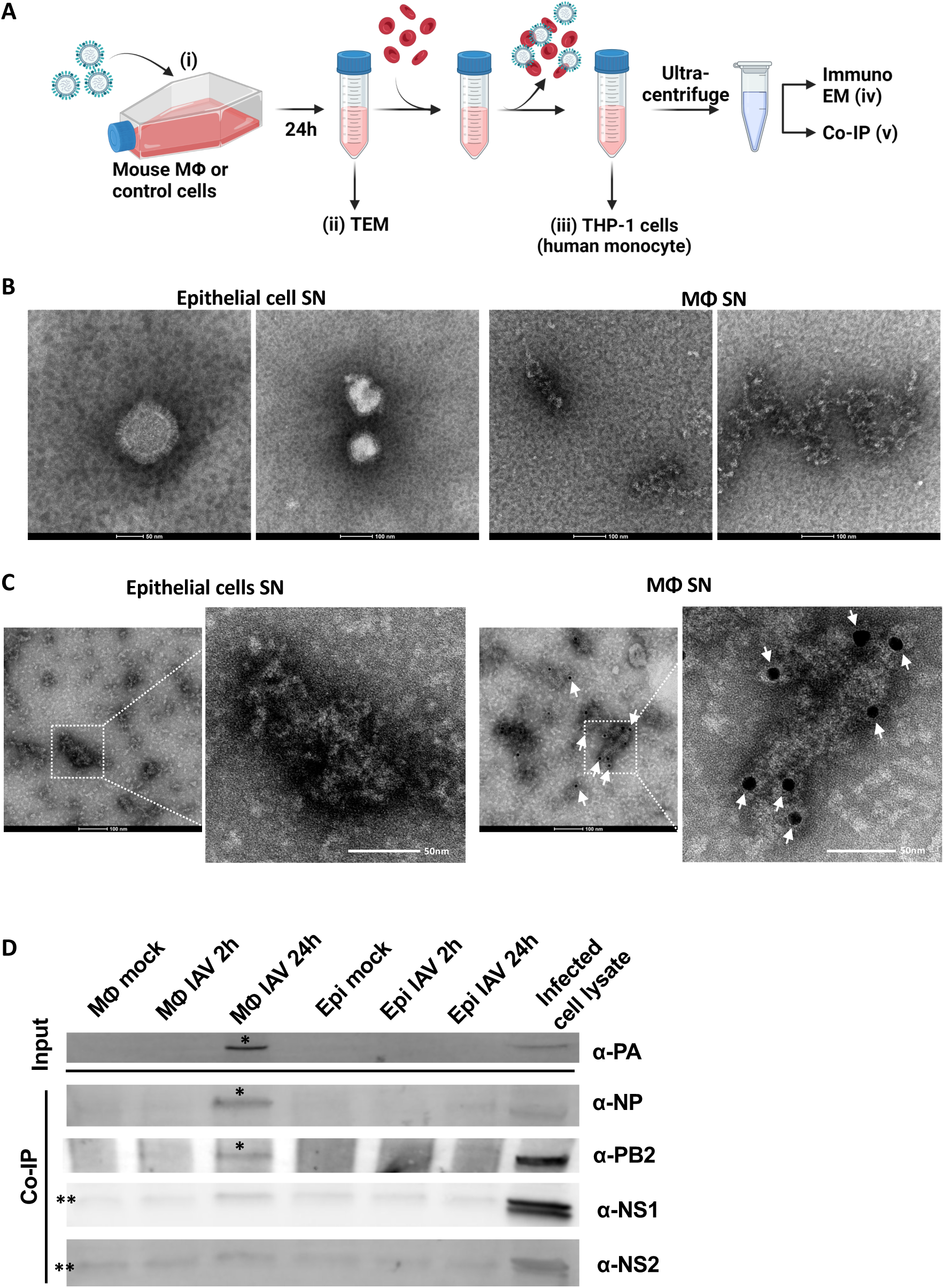
vRNA is released from IAV infected macrophages as packaged vRNP complexes. (A) Schematic overview of experimental design to remove residual input virions from SN of IAV infected MΦ following abortive replication, as well as newly synthesised infectious viral particles released from epithelial cells during productive replication. MΦ and epithelial cells were infected with IAV (MOI 5), or mock infected, and cell-free SNs were collected at 2 and 24 hpi. Depletion of virions was then achieved by addition of turkey RBCs, followed by centrifugation to pellet the cells and associated virus. vRNP complexes in virus-cleared SNs were concentrated by ultracentrifugation. **(B)** Epithelial cells (MDCK) and MΦ (RAW264.7) were infected with IAV (MOI 5) and cell culture SN was negative stained and analysed 24 hpi via TEM ((A) dashed box). SN samples were processed as described in **(A)** and analysed via **(C)** TEM or **(D)** co-immunoprecipitation. **(C)** SN samples of MΦ and epithelial control cells were stained with anti-NP antibody and immunogold labelled. Arrow heads show gold particles in MΦ SN only. **(D)** Protein A-Sepharose beads coated with anti-PA antibody were added to SN samples (mock infected, 2 and 24 hpi) from both cell types and incubated overnight. PA and any associated viral proteins were then co-immunoprecipitated and analysed via immunoblotting. The presence of individual IAV vRNP proteins was detected using PA (83 kDa), NP (56 kDa) and PB2 (86 kDa) specific antibodies (*indicates presence of PA, NP and PB2 proteins as vRNP complexes in IAV-infected MΦ SN and not epithelial SN). Antibodies against non-structural IAV proteins (NS1 (26 kDa) and NS2 (14 kDa)) were used as specificity controls (** indicates a non-specific band for NS1 and NS2 present in all samples including uninfected cell SN). Whole-cell lysates of IAV infected MΦ (infected cell lysate) was used as a positive control to show expression of each of the aforementioned IAV proteins.

### vRNP complexes released from IAV-infected MΦ induce IL-1β expression in uninfected cells

We next explored if vRNP complexes released from IAV-infected mouse MΦ were able to stimulate inflammatory or antiviral responses in uninfected cells. To this end, we exposed human THP-1 cells to vRNP complexes present in virus-depleted SN from IAV-infected mouse MΦ and epithelial cells (Fig. 2A (iii)). Human cells were chosen to reduce the impact of mouse cytokines present in the IAV-infected MФ and epithelial cell SNs. We measured mRNA expression of selected antiviral (IFNβ, Mx1 and IFITM1) and inflammatory genes (TNFα, IL-1β, IL-6) at 6 hours post exposure by RT-qPCR (Fig. 3A/B). Recombinant human IFNα, a RIG-I agonist (ppp-dsRNA) and a TLR7/8 agonist (R848) were used as controls to stimulate antiviral and inflammatory host responses via specific pattern recognition receptors (PRRs). We observed no significant changes in mRNA expression for any of the selected antiviral genes, including IFNβ (Fig. 3A (i-iii)), when uninfected cells were exposed to vRNPs present in IAV-infected MΦ SN or to control SN from IAV-infected epithelial cells, despite robust induction by control agonists including ppp-dsRNA (Fig. 3A (i-iii)). Similarly, the induction of TNFα was modest and no significant induction of IL-6 was observed when uninfected cells were exposed to MΦ SN containing vRNPs (Fig. 3B (i/ii)). In contrast, mRNA expression of inflammatory IL-1β was significantly increased in uninfected cells exposed to SN containing vRNP from IAV-infected MΦ, but not by SN from IAV-infected epithelial cells or uninfected MΦ (Fig. 3B (iii)). Inflammatory responses were also induced in an alternative human cell line (A549) exposed to vRNP complexes isolated from IAV-infected mouse MΦ, where significant increases in both IL-1β and IL-6 mRNA were observed (data not shown). As, IL-1β expression was also significantly increased in uninfected THP-1 cells exposed to RIG-I agonist (>45x) or TLR7/8 agonist (>200x) (Fig. 3B (iii)), we tested if the vRNPs present in SN of IAV-infected MΦ might be specifically sensed through either the RIG-I or TLR7/8 PRRs. For this, we exposed uninfected THP-1 cells to a TLR7/8 antagonist prior to exposure to SN containing vRNPs from IAV-infected MΦ (Fig. 3C). While THP-1 cells treated with the TLR7/8 antagonist prior to stimulation with the TLR7/8 agonist showed reduced IL-1β induction, stimulation with SN containing vRNPs from IAV-infected MΦ did not display reduced IL-1β induction (Fig. 3C). This indicates that induction of IL-1β expression through vRNPs is unlikely to be mediated through TLR7/8 in THP-1 cells. Next, THP-1 cells deficient in mitochondrial antiviral-signalling protein (MAVS^-/^) or deficient in both retinoic acid-inducible gene I and melanoma differentiation-associated protein (RIG-I/MDA5^-/-^) were exposed to SN containing vRNPs from IAV-infected MΦ, and both displayed significantly less IL-1β induction when compared to wildtype cells (Fig. 3D). We confirmed that both WT and KO THP-1 cells showed similar levels of induction when treated with R848 (data not shown). Although IL-1β mRNA induction was reduced in both MAVS^-/-^ and RIG-I/MDA5^-/-^ THP-1 cells, it was not completely inhibited, suggesting IAV vRNP complexes may be sensed through alternative receptors or IL-1β mRNA is also induced through other signalling molecules in the SN of infected MΦ.

**Figure 3:**
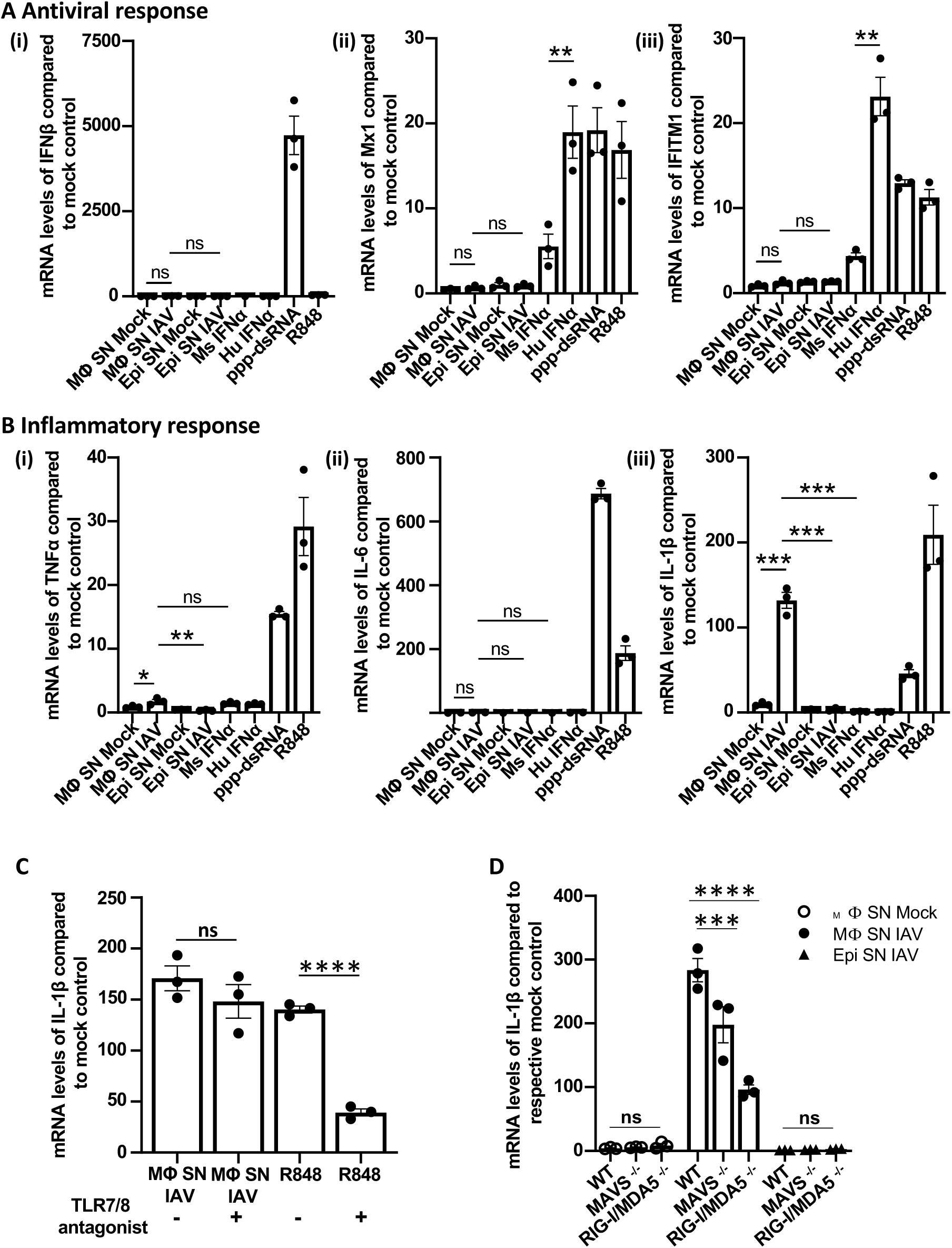
vRNP complexes isolated from IAV-infected mouse MΦ induce IL-1β expression in uninfected cells (A-D) Human monocytes (THP-1) were exposed to IAV vRNP complexes in virus-cleared (see Fig. 2A) cell-free SN from infected (24h) or uninfected (mock) mouse MΦ or epithelial cells, or stimulated with mouse IFNα, human IFNα, RIG-I agonist (ppp-dsRNA in complex with Lipofectamine®) or TLR7/8 agonist (R848) for 6h. mRNA expression of **(A)** IFNβ, **(B)** Mx1, **(C)** IFITM1 or **(D)** IL-1β was assessed via RT-qPCR and fold induction compared to untreated cells was calculated. **(E)** THP-1 cells were treated with TLR7/8 antagonist (ODN20959) or left untreated before cells were either stimulated with R848 or the virus-cleared SN of mouse MΦ infected for 24h. Induction of IL-1β mRNA compared to untreated cells was analysed via RT-qPCR. **(F)** THP-1 WT, MAVS^-/-^ and RIG-I/MDA5^-/-^ cells were treated with virus-cleared SN from uninfected and infected MΦ and infected epithelial cells for 6h and analysed for the expression of IL-1β mRNA via RT-qPCR. Representative experiments of n=3 are shown. ANOVA or t-test was performed with p<0.5 *, p<0.1 **, p<0.01 ***, p<0.001 ****, ns: not significant.

### vRNP-containing MΦ SN triggers a global inflammatory response in THP-1 cells

To gain further insights into the breadth of antiviral vs inflammatory responses, we performed RNAseq on uninfected human cells exposed to the vRNP complexes from IAV-infected mouse MΦ SN (Fig. 4). THP-1 cells exposed to vRNP complexes in SN from IAV-infected mouse MΦ showed upregulation of 122 genes and downregulation of 8 genes (compared to those exposed to SN from uninfected MΦ) and upregulation of 322 genes and downregulation of 56 genes (compared to those exposed tp IAV-infected epithelial SN). (Fig. 4A). Interestingly, there were more differentially up/down-regulated genes observed when comparing THP-1 cells exposed to SN from infected epithelial cells and infected MΦ than when comparing THP-1 cells exposed to SN from uninfected MΦ and infected MΦ, highlighting the differences in steady-state gene expression of MΦ compared to epithelial cells. Upregulated genes in THP-1 cells stimulated with vRNPs from IAV-infected MΦ compared to IAV-infected epithelial cell SN or mock-infected MΦ SN were associated with GO terms for immune response, chemokine-mediated signalling and inflammatory response (Fig. 4B). Most upregulated genes were associated with inflammation and immune response (Fig. 4D) and in addition to IL-1β (4.6 logFC), expression of signalling genes such as CXCL13 (8.7 logFC), IL-32 (6.5 logFC), CCL4 (5.4 logFC) and CXCL10 (4 logFC) were amongst the most significantly upregulated (Fig. 4C). Similar to our RT-qPCR analysis on antiviral genes (IFNβ, Mx1, IFITM1, Fig.3) only a few genes associated with ‘response to virus’ GO terms like OASL (2.2 logFC) or IRF7 (1.7 logFC) or were elevated in THP-1 cells exposed to vRNP complexes in SN from IAV-infected mouse MΦ (Fig. 4E). Together, these data demonstrate that IAV vRNPs released from infected mouse MΦ during abortive replication have the potential to shape the inflammatory and cytokine/chemokine response in neighbouring uninfected cells. This provides novel insights into the functional consequences of IAV abortive replication in MΦ.

**Figure 4:**
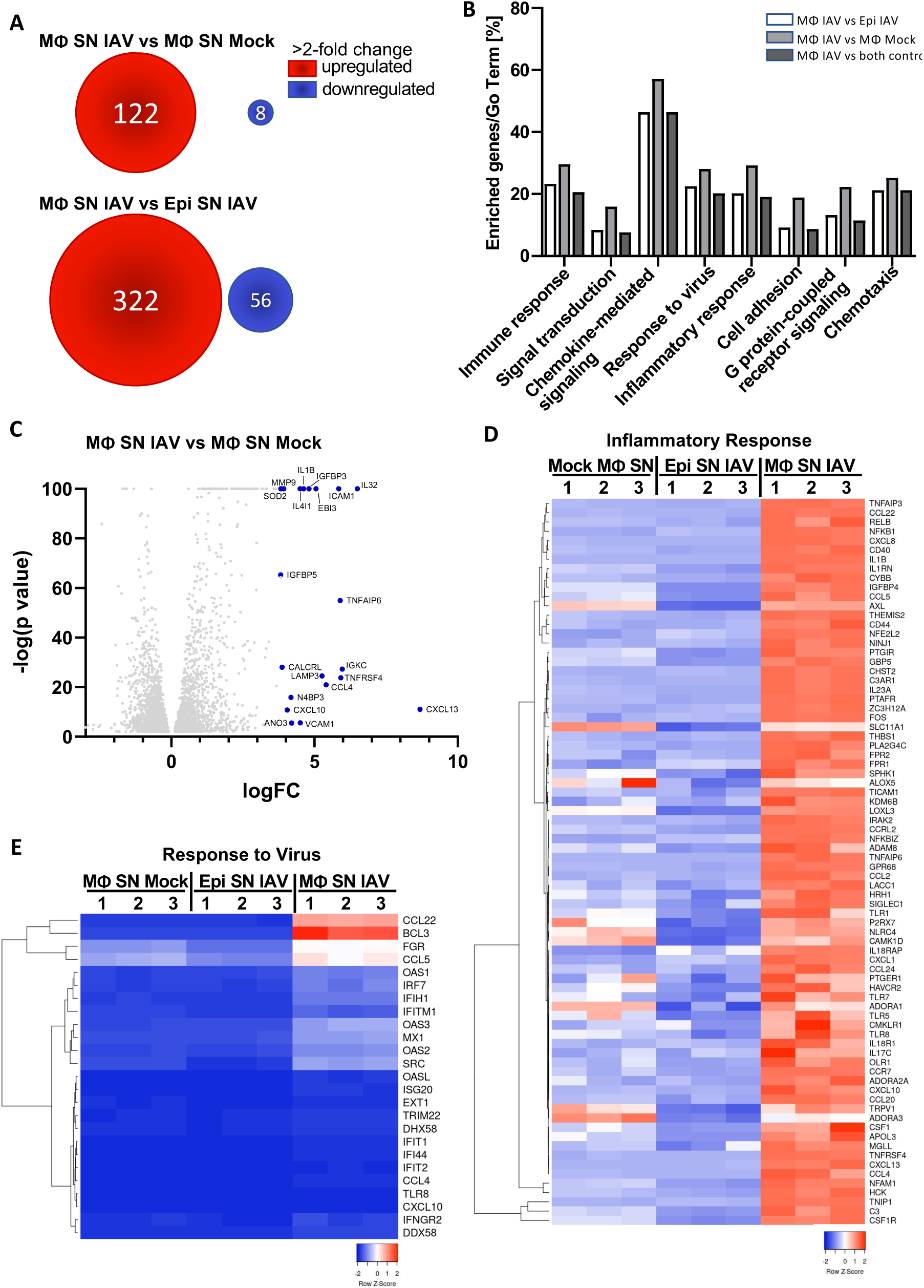
vRNP-containing MΦ SN triggers a global inflammatory response in THP-1 cells. THP-1 cells were exposed to vRNP in virus-cleared SN from IAV infected MΦ or to virus-cleared SN from IAV infected epithelial cells or to SN from uninfected MΦ for 6h. Cellular mRNA levels were assessed through Illumina RNA sequencing **(A)** Total number of genes up- or downregulated in THP-1 cells exposed to virus-cleared SN from infected MΦ compared to cells exposed to SN from mock infected MΦ or infected epithelial cells. Number of genes with fold change >2 is shown. **(B)** Genes expressed in THP-1 cells exposed to infected MΦ SN and associated with enriched GO-terms (%) compared to infected epithelial cells or uninfected MΦ is shown. **(C)** Comparative expression of genes in THP-1 cells exposed to IAV MΦ SN compared to Mock MΦ SN (log FC, x-axis). -log(p value) is shown on y-axis and the 20 most differentially expressed genes are highlighted in blue. **(D)** Relative expression of selected host genes associated with ‘*<u>immune response and inflammation’</u>* comparing THP-1 cells exposed to Mock MΦ SN, virus-cleared IAV epithelial cell SN and vRNPs in virus-cleared IAV MΦ SN. Downregulated genes are represented in blue while upregulated genes are presented in red. **(E)** Relative expression of host genes associated with *‘<u>Response to Virus’</u>* GO term in THP-1 cells exposed to Mock MΦ SN, virus-cleared IAV Epithelial cell SN and vRNPs in virus-cleared IAV MΦ SN. Downregulated genes are represented in blue while upregulated genes are presented in red. 1, 2, 3 represent triplicate samples.

## Discussion

We have previously shown that infection of mouse MΦ by seasonal IAV leads to abortive replication, where viral RNA is transcribed and viral proteins are translated within the cell, but infectious viral particles are not released ^18^. Surprisingly, herein we detected significant amounts of viral RNA and NP present in the SN of IAV-infected MΦ during abortive replication (Fig. 1, Fig S2). EM analysis of the SN showed the presence of vRNP-like complexes and immunogold labelling indicated that these were associated with IAV NP (Fig. 2A). Furthermore, IAV PA released from IAV infected MΦ was in complex with PB2 and NP (Fig. 2B), suggesting that vRNPs are released during abortive replication. We performed NGS analysis to show that all RNA segments were present in the SN of infected MΦ during abortive replication (Fig. 1C). Even though we observed slight differences in the abundance of some RNA segments (e.g. NA), we could detect significant amounts of each RNA segment in MΦ SN, suggesting that deficient production of specific viral RNA segments is not the cause of abortive replication in MΦ.

Following removal of any residual virions from the initial infection, we show that vRNP complexes in SN from IAV-infected mouse MΦ are potent viral PAMPs that can be sensed at least in part through the RIG-I/ MDA5 pathway in uninfected human THP-1 cells resulting in the induction of IL-1β transcription (Fig. 3). We observed a significant reduction of IL-1β mRNA levels in RIG-I/MDA5^-/-^ and MAVS^-/-^ THP-1 cells when exposed to the vRNP-containing MΦ SN, while TLR7/8 antagonists had no effect. Although the RIG-I/MDA5 pathway seems important for driving the IL-1β response, it was not completely abolished in either the MAVS^-/-^ nor in the RIG-I/MDA5^-/-^ THP-1 cells. This could be due to the vRNP complexes being sensed through other external or internal sensors. Indeed a recent study in epithelial cells showed that IAV NP, which is part of the vRNP complex can be recognised by TLR2 and TLR4 on the cell surface ^23^. Considering the high levels of NP detected in the SN of infected MΦ (Fig 1 D), these receptors might be engaged in the response to vRNPs. We cannot exclude that cross-reactive mouse cytokines and chemokines present along with vRNP complexes in IAV-infected MΦ SN contribute to the inflammatory response in uninfected human THP-1 cells, but several observations suggest that this is unlikely. Firstly, mouse cytokines and chemokines were also detected in SN from control IAV-infected mouse epithelial cells (Fig Sl. 3) but did not induce significant inflammatory responses in THP-1 cells (Fig. 3B). Secondly, exposure of THP-1 cells to recombinant mouse TNFα and IL-1β at levels detected in SN from IAV infected MΦ (3ng/mL and 30pg/mL, respectively) did not induce significant IL-1β mRNA expression (SI 3B). Therefore, we propose that significant levels of vRNA and viral protein (NP) released by MΦ during IAV abortive replication present potent PAMPs that influence the immune response to IAV infection in the local microenvironment. When RNA sequencing was used to examine the global response of uninfected cells exposed to the SN of infected MΦ, we observed a strong inflammatory response with the upregulation of cytokine and chemokine gene expression and inflammatory gene expression (Fig. 4). In contrast, we observed very few antiviral genes or genes associated with ‘Response to Virus’ GO terms that were upregulated in human THP-1 cells exposed to vRNP-containing SN from mouse MΦ. Instead, uninfected cells responded with a general inflammatory response, which shapes the infectious milieu and both the innate and adaptive response by attracting distinct immune cells. CXCL13, which was among the most highly expressed genes in THP-1 cells exposed to SN from infected MΦ, is essential for attracting B-lymphocytes ^24^, whose migration to the site of infection is essential for clearance of IAV infection ^25,26^. IL-32 was another significantly induced host factor in the THP-1 cells and has antiviral properties against IAV and other viruses like hepatitis B virus ^27,28^. In the presence of IL-32, IAV genome replication and mRNA synthesis are greatly reduced, and viral replication is heavily impaired ^27^.

While the protective role of MΦ during IAV infection may in part be due to limited release of new viral progeny through abortive replication ^17,18^, we now propose that vRNP complexes released from IAV-infected MΦ have the potential to shape local inflammatory responses in uninfected cells. Our findings give new insight into MΦ responses during respiratory virus infection. Future studies are required to define the balance of MΦ-driven pathogenic inflammatory responses vs anti-viral defense in the context of influenza disease.

## Acknowledgments

This work was supported by Project Grant APP1184532 from the National Health and Medical Research Council (NHMRC) of Australia. The Melbourne WHO Collaborating Centre for Reference and Research on Influenza is supported by the Australian Government Department of Health. We thank the Melbourne Flow Cytometry Core Platform for assistance with flow cytometric analysis. We thank Marcel Doerflinger from the Walter and Eliza Hall Institute (Melbourne, Australia) for generously providing the RIPK3 inhibitor and Thomas Zillinger from the University of Bonn for providing the THP-1 MAVS^-/-^ and RIG-I/MDA5^-/-^ cells. Josh Zhang at the University of Melbourne kindly assisted with curation of the RNASeq data.

## Author contributions

Conceptualization, S.F., S.L.L., P.C.R.; Methodology, S.F., S.L.L., P.C.R.; Formal Analysis, S.F., V.L.S, J.A.R., L.J.C., Y.M.D., J.M., S.L.L.; Investigation, S.F., V.L.S, J.A.R., Y.M.D., J.M., S.L.L.; Writing – Original Draft, S.F., S.L.L.; Writing – Review & Editing, S.F., S.L.L., L.I.L, S.S.L., P.C.R. A.G.B.; Resources, J.A.R., J.M.M. P.C.R.; Visualization, S.F., J.A.R., L.J.C; Project Administration and Supervision, S.L.L.; Funding acquisition, S.L.L., P.C.R., A.G.B.

## Methods

### Cell lines and primary cells

Mouse MΦ cell line RAW264.7 (American type culture collection (ATCC), Manassas, VA) was grown in DMEM media containing 10% FCS (Bovogen Biologicals), 2mM GlutaMAX™ and 1mM sodium pyruvate (all Gibco, Thermo Fisher Scientific, MA, USA). Madin-Darby Canine Kidney (MDCK) cells and human THP-1 cells (ATCC) were maintained in RPMI media with 10% FCS, 2mM GlutaMAX™ and 1mM sodium pyruvate (all Gibco). THP-1 RIG-I/MDA5^-/-^ and MAVS^-/-^ KO cell lines were provided by Thomas Zillinger (University of Bonn, Germany) and cultured in RPMI media with 10% FCS, 2mM GlutaMAX™ and 1mM sodium pyruvate (all Gibco). Murine lung epithelial LA-4 cells (ATCC) were cultured in Kaighn’s modification of Ham’s F-12 medium supplemented with 10% FCS, 2mM GlutaMAX™ and 1mM sodium pyruvate (all Gibco). Resident peritoneal exudate cell (PEC) mouse MΦ were obtained from C57BL/6 mice as previously described (REF) and grown in DMEM media containing 10% FCS, 2mM GlutaMAX™ and 1mM sodium pyruvate (all Gibco). Mice were maintained in a pathogen-free environment at the Doherty Institute (University of Melbourne, Australia) and used at 6-10 weeks of age. All experiments were approved and conducted according to the guidelines of the University of Melbourne Animal Ethics Committee.

### Influenza A virus and infection assays

Influenza A virus strain BJx109, a high-yielding reassortant of A/Beijing/353/89 (H3N2) with A/PR8/34 bearing the H3N2 surface glycoproteins, was used in this study as allantoic fluid propagated in embryonated hens eggs at the WHO Collaborating Centre for Reference and Research on Influenza (WHO CCRRI), Melbourne, Australia (ethics approval from the University of Melbourne Biochemistry & Molecular Biology, Dental Science, Medicine, Microbiology & Immunology, and Surgery Animal Ethics Committee). For infection assays, cells were seeded the day prior to the infection, washed 2x in FCS-free media containing supplements (2mM GlutaMAX™, 1mM sodium pyruvate) and infected with BJx109 at the indicated MOI in a low volume. Infections were performed at 37°C for 1h. The inoculum was removed, and cells were washed 2x in FCS-free media with supplements. Cells were maintained in FCS-free media containing supplements for the duration of the infection.

### Plaque assay for quantitation of infectious viral particles

MCDK cells were seeded in 6-well plates at a high density to achieve 90-95% confluency on the day of the assay. In the absence of trypsin in the samples to be tested, virus was activated by adding 8μg/mL trypsin (TPCK treated, Worthington, Lakewood, NJ) in FCS-free media to samples in a 1:1 ratio. Samples were incubated for 30 min at 37°C and 1/10 serial dilutions were set up. Cells were washed in FCS-free media and inoculated with 150uL of virus dilution. Infections were performed at 37°C for 1h with regular agitation. Overlay (L15 media (Gibco) with 0.8mM HEPES, penicillin/streptomycin (both Gibco), sodium carbonate (Sigma) and 0.9% agarose) was added to the cells in two steps and allowed to cool. Cells were incubated for 3 days at 37°C and the formation of plaques was monitored. Based on the initial inoculum (150uL) plaque forming units (PFU) per mL were calculated.

### RT-qPCR for host and viral genes

RNA was extracted from cells using the RNAeasy Plus Kit (QIAGEN), while viral RNA present in cell culture SN was isolated using the QIAamp Viral RNA Mini Kit (QIAGEN). RNA was DNAse treated and cellular RNA samples were adjusted to the same concentration. For vRNA analysis, cDNA was generated using the Tetro cDNA Synthesis Kit (Bioline) and the Uni12 primer (5’-AGCAAAAGCAGG-3’, Sigma). For mRNA analysis Oligo(dT) primers were used to create the cDNA. RT-qPCR was performed using SensiFAST™ SYBR® Lo-ROX Kit (Bioline) and either the QuantStudio™ 5 or QuantStudio™ Flex 7 analyser (Applied Biosystems). For analysis on viral gene copy numbers, gene-specific plasmid standards with a known concentration and copy number were examined alongside the samples. To analyse the expression of host genes, housekeeping genes and untreated controls were used to calculate the fold expression via the ΔΔCt method. The following primer sets were used: M (F: 5’-GACCRATCCTGTCACCTCTGAC-3’, R: 5’-GGGCATTYTGGACAAAKCGTCTACG-3’), NA (F: 5’-CAACCAAGTAATGCCGTGTG-3’, R: 5’-TTGTCACCCAAATGTCTCCA-3’), PA (F: 5’-ACAAGGCATGCGAACTGACA-3’, R: 5’-TGGAGCCACATCTTCTCCAA-3’), PB1 (F: 5’-CTCTGACGATTTTGCTCTGATTGT-3’, R: 5’-TCGACTCCGGCTTGAATCC-3’). HPRT (F: 5’-TCAGGCAGTATAATCCAAAGATGGT-3’, R: 5’-AGTCTGGCTTATATCCAACACTTCG-3’), IFNβ (F: 5’-CAGTCCTGGAAGAAAAACTGGAGA-3’, R: 5’-TTGGCCTTCAGGTAATGCAGAA-3’), Mx1 (F: 5’-GTTTCCGAAGTGGACATCGCA-3’, R: 5’-CTGCACAGGTTGTTCTCAGC -3’), IFITM1 (F: 5’-AGCATTCGCCTACTCCGTGAAG-3’, R: 5’-CACAGAGCCGAATACCAGTAACAG -3’), TNFα (F: 5’-TGCCTGCTGCACTTTGGAGTGA-3’, R: 5’-AGATGATCTGACTGCCTGGGCCAG-3’), IL-6 (F: 5’-CTCAGCCCTGAGAAAGGAGACAT-3’, R: 5’-TCAGCCATCTTTGGAAGGTTCA-3’), IL-1β (F: 5’-CTGATGGCCCTAAACAGATGAAGT-3’, R: 5’-AGCCCTTGCTGTAGTGGTGGT-3’).

### AlphaLISA® assay for IAV NP expression

AlphaLISA® detection kit targeted against IAV NP protein was purchased from Perkin Elmer and performed according to the manufacturers protocol. In brief, 5μL of sample or NP standard were added to 96-well half area plates (Perkin Elmer) and AlphaLISA anti-NP acceptor beads (10 μg/mL final concentration) were added and incubated for 30 min at room temperature (RT). Biotinylated anti-NP antibody (2 nM final concentration) was added and incubated for 60 min at RT. Lastly, samples were combined with SA-donor beads (40 μg/mL final concentration) and incubated for 30 min at RT and protected from light. The assay was measured with an EnVison-Alpha plate reader (Perkin Elmer) and analysed via the standard curve.

### Slot blot analysis of IAV NP expression

Cell culture media was analysed for the presence of IAV NP by concentrating the sample on a PVDF membrane. For this either neat SN or 10-fold dilutions were applied to the slots of a slot-blot manifold (Masterflex, Radnor, US) and transferred to the PVDF membrane by applying vacuum pressure. Membranes were blocked in 5% BSA for 60min at RT and incubated with the primary antibody (anti-NP, Bio-Rad, 1/5000) for 45-60min shaking. The membrane was washed in phosphate-buffered saline (PBS) with 0.05 % Tween®20 (Sigma) and the secondary antibody was added for 30min (anti-mouse-HRP, Dako, 1/10000) shaking. Membranes were first washed in PBS-T and then in PBS only, before TrueBlue Peroxidase Substrate (SeraCare, Milford, US) was added to visualise the bound NP antibody.

### Next generation sequencing (NGS) of viral gene segments

RNA was extracted from 140 µl of cultured media using QIAamp RNA extraction kit (Qiagen) according to manufacturer’s instruction. Multi RT-PCR was performed using universal influenza A primers to amplify all eight segments as described ^29^, briefly two primers (Uni13/Inf-1 and Uni12/Inf-1) were used with SuperScript III One-step RT-PCR with Platinum Taq High Fidelity kit as per instruction. A 50 µl reaction with 5 µl of RNA and 1 µl of each primer (20uM stock) was set up for each sample for RT-PCR using the following program: 55°C - 60 min, 94°C - 2 min, 5 cycles of 94°C – 30 sec, 45°C – 30 sec, 68°C -4 min, 31 cycles of 94 °C - 30 sec, 57 °C - 30 sec, 68 °C - 4 min, 68 °C - 5 min and hold at 4°C. Amplicons were analysed on TapeStation 4200 (Invitrogen) to determine the quality and quantity. 200 ng of PCR product for each sample were normalized in 30 µl volume for NGS library construction using Illumina DNA Prep (M) Tagmentation kit (Illumina). IDT for Illumina Nextera DNA Unique Dual Indexes were used for pooling multiple samples together, pooled library was quantified using Tapescreen HSD1000 on TapeStation 4200, then diluted to 150 pM in elution buffer supplied in the library prep kit and loaded to the iSeq100 flow cell for sequencing on iSeq100 (Illumina). After the NGS run, fastq data were retrieved and analysed using IRMA pipeline ^30^ to generate consensus sequences and genome coverage diagram, the coverage diagrams were used for defective interfering (DI) particle investigation. The sequences for all eight influenza A genes were identical to the parent virus sequence.

### Virus-depletion and concentration of IAV cell culture SN containing vRNPs

Cells were either infected with IAV (MOI 5) or mock infected and cell culture SN was collected at 2 and 24 hpi. Cell debris was removed by centrifugation at 1800 rpm for 5 min. For each sample 10 mL of a 5 % turkey red blood cells (kindly supplied by the WHO Collaborating Centre for Reference and Research on Influenza, Melbourne, Australia) suspension was centrifuged at 1500 rpm for 3 min and SN was discarded. RBCs were resuspended in cell-free culture medium of mock/infected cells and incubated for 2h at 4°C rotating. RBCs were removed through centrifugation and cell culture SN was transferred to a new tube containing fresh RBCs (sedimented from 10 mL of 5 % RBC suspension). RBCs and SN were again incubated for 2 h at 4 °C rotating. Cell culture SN was centrifuged at 1500 rpm for 3 min 2x to remove any residual RBCs and transferred to a new tube. For functional readout experiments, small aliquots were stored at -80 °C. For TEM and co-immunoprecipitation analysis, virus-depleted SNs were centrifuged at 28,000 rpm at 4 °C for 2 h. SN was removed and samples were resuspended in 200 μL PBS.

### TEM and immunogold labelling of vRNPs

For negative staining, 6 μL of supernatant was applied directly to glow-discharged, 400-mesh copper formvar-carbon coated grids (EMgrid) and allowed to adsorb for 30 seconds. Supernatant was removed by blotting with a wedge of Whatman number 1 filter paper, and then negative stained for 10 seconds using 1% aqueous uranyl acetate (Electron Microscopy Supplies) and blotted to remove excess stain.

Indirect immuno-gold electron microscopy was performed by suspending glow-discharged grids on a 10µL drop of supernatant for 5 minutes, then washed three-times in HN buffer, consisting of 10mM HEPES in saline (pH 7.4). Washed grids were then suspended on a 25 µL drop of HN buffer containing 1% bovine serum albumin (BSA) and incubated in a room-temperature humidity chamber for 20 minutes. Grids were then blotted dry and suspended on a 25 µL drop of primary anti-NP antibody (Bio-Rad), diluted 1:500 in HN diluent, consisting of HN buffer and 0.2% BSA. The grids were then incubated for 1 hour in a humidity chamber at room temperature, washed tree times in HN diluent, then suspended on a 25 µL drop of 10nm gold-conjugated anti-Mouse IgG (Sigma-Aldrich), at a dilution of 1:20 in HN diluent. Grids underwent a final incubation in a room temperature humidity chamber for 1 hour, washed five times in HN diluent, then rinsed three times in 0.22µm filtered MilliQ water. The grids were then negative stained using 1% aqueous uranyl acetate, then blotted to remove excess stain.

Negative stained samples were air-dried for 10 minutes then examined using an FEI Tecnai T12 Spirit electron microscope operating at an acceleration voltage of 80 kV. Final images were collected using an FEI Eagle 4k CCD camera at native resolution.

### Co-Immunoprecipitation of vRNPs

All co-immunoprecipitation steps were carried out at 4 °C unless otherwise specified. Protein A-Sepharose™ beads were blocked in 0.1 % BSA for 1 h and then coated with anti-PA antibody (Pierce) for 3h rotating. Virus-depleted and concentrated cell SN (in PBS) was added to anti-PA (Thermo Fisher) coated beads and incubated overnight. Beads were washed 6x in PBS to remove any residual unbound proteins. Lysis buffer (50 mM Tris-HCl (pH 7.5), 150 mM NaCl, 0.5% (vol/vol) Triton X-100, 1 mM CaCl2, 1 mM MgCl2) containing protease inhibitors (Roche, Germany) was added to beads and incubated for 10 min to ensure lysis. SDS loading dye (300 mM Tris-HCl pH 6.8, 10% Sodium dodecyl sulphate (SDS), 30% glycerol, 0.02% Bromophenol blue) was added to each sample and proteins were detached from beads through incubation at 95 °C for 10 min. Beads were centrifuged, and eluted proteins were transferred to a new tube until analysis via immunoblotting.

### Immunoblot analysis

Cells were washed in PBS prior to lysis in lysis buffer containing protease inhibitors. Lysates were incubated for 30 min at 4°C under agitation and spun at 13,000 rpm for 10 min at 4°C. Clarified lysates were transferred to a new tube and SDS loading dye was added. Samples were heated to 95°C for 5 min and separated on 10-15% SDS-PAGE gels (depending on protein size). Proteins were transferred to PVDF membrane (Millipore) and blocked in 5% bovine serum albumin (BSA, Sigma) and 0.05 % Tween®20 for at least 2h. Membranes were stained with primary antibody in blocking buffer at 4°C overnight: anti-NP (1:5000, OBT1555, Bio-Rad), anti-PA, anti-PB2, anti-NS1 and anti-NS2 (all 1:2000, Thermo Fisher). Membranes were washed in PBS-T and incubated with fluorescent secondary antibody (anti-mouse or anti-rabbit conjugated to AlexaFluor488 or 647, both Life Technologies) for 2-3h at RT. After several wash steps in PBS-T, membranes were washed in PBS before imaging on the AI600 imager (Amersham Biosciences).

### Immunofluorescence microscopy

Cells were seeded on glass coverslips, treated according to the experiment and fixed in 4% paraformaldehyde (PFA, Electron Microscopy Sciences) for 15min at RT. If samples were stained on the surface, cells were incubated with primary antibody in blocking buffer first, before cells were fixed and permeabilised. Samples were washed in PBS and permeabilised with 0.1% Triton™ X-100 (Sigma) in PBS for 15min at RT. Background staining was reduced by blocking samples in 5% BSA and 5% FCS in PBS for 60min at RT. Primary antibody was added in blocking buffer for 45-60min at the indicated concentration: anti-NP (1:50, clone MP3.10g2.1C7) and anti-HA (1:500) were obtained from the WHO Collaborating Centre for Reference and Research on Influenza, Melbourne, Australia), PA (1:250), PB2 (1:250), NS1 (1:250, PA5-32243 and NS2 (1:250, PA5-32234) were all purchased from Thermo Scientific, Rockford, USA, M2 (1:200). After washing in PBS, samples were incubated with secondary antibodies conjugated to AlexaFluor 488, 568 or 647 (all from Life Technologies) for 30-45min. Finally, coverslips were washed in PBS followed by Milli-Q® H2O and dried before mounting with ProLong™ Diamond Antifade Mountant (Life Technologies). Images were acquired using the Zeiss LSM780 confocal microscope (63x objective) and analysed using ImageJ software.

### Assessing the immunoreactivity of vRNPs on uninfected cells

RT-PCR of selected host genes: Human THP-1 cells in suspension were either left untreated, induced with mouse IFNα (1000 units/mL, Miltenyi Biotech), human IFNα (1000 units/mL, Miltenyi Biotech), RIG-I agonist (ppp-dsRNA in complex with Lipofectamine®, Thermo Fisher), TLR7/8 agonist (R848, 8 μg/mL, InvivoGen) or exposed to vRNPs in virus-cleared SN from infected (24 hpi) or uninfected MΦ (RAW 264.7) and epithelial cells (LA4). All treatment groups were supplemented with 1 % FCS and RNA was extracted in RNAeasy Plus lysis buffer (QIAGEN) 6 h post treatment.

RNA sequencing: THP-1 cells were exposed to vRNPs in virus-cleared SN from infected mouse MΦ, infected mouse epithelial cells (both 24 hpi) and uninfected mouse MΦ as described above. RNA was extracted using RNeasy Plus mini kit (QIAGEN). Residual DNA was digested with RNAse-free DNAse for 30 min (Promega), before enzyme and buffer were removed via column clean up (QIAGEN). RNA was eluted in 20μL RNAse-free water and processed for Illumina mRNA sequencing through the Australian Genome Research Facility (AGRF).

RNA sequencing analysis: Kallisto ^31^ (quant subcommand, version 0.44.0) was used to generate gene counts for all sequenced samples, using the reference transcriptome GRCm38.96. Subsequently, all analysis was carried out using the R language (version 4.0.5). The tximport library was used to read Kallisto output, and the DESeq2 package ^32^ was used calculate differential expression p-values and log fold ratios between conditions. False discovery rates were calculated using the Benjamini Hochberg correction.

