## Supplemental figures for "Influenza infected macrophages release viral ribonucleoproteins that shape the local host response"

Suppl. Figure 1

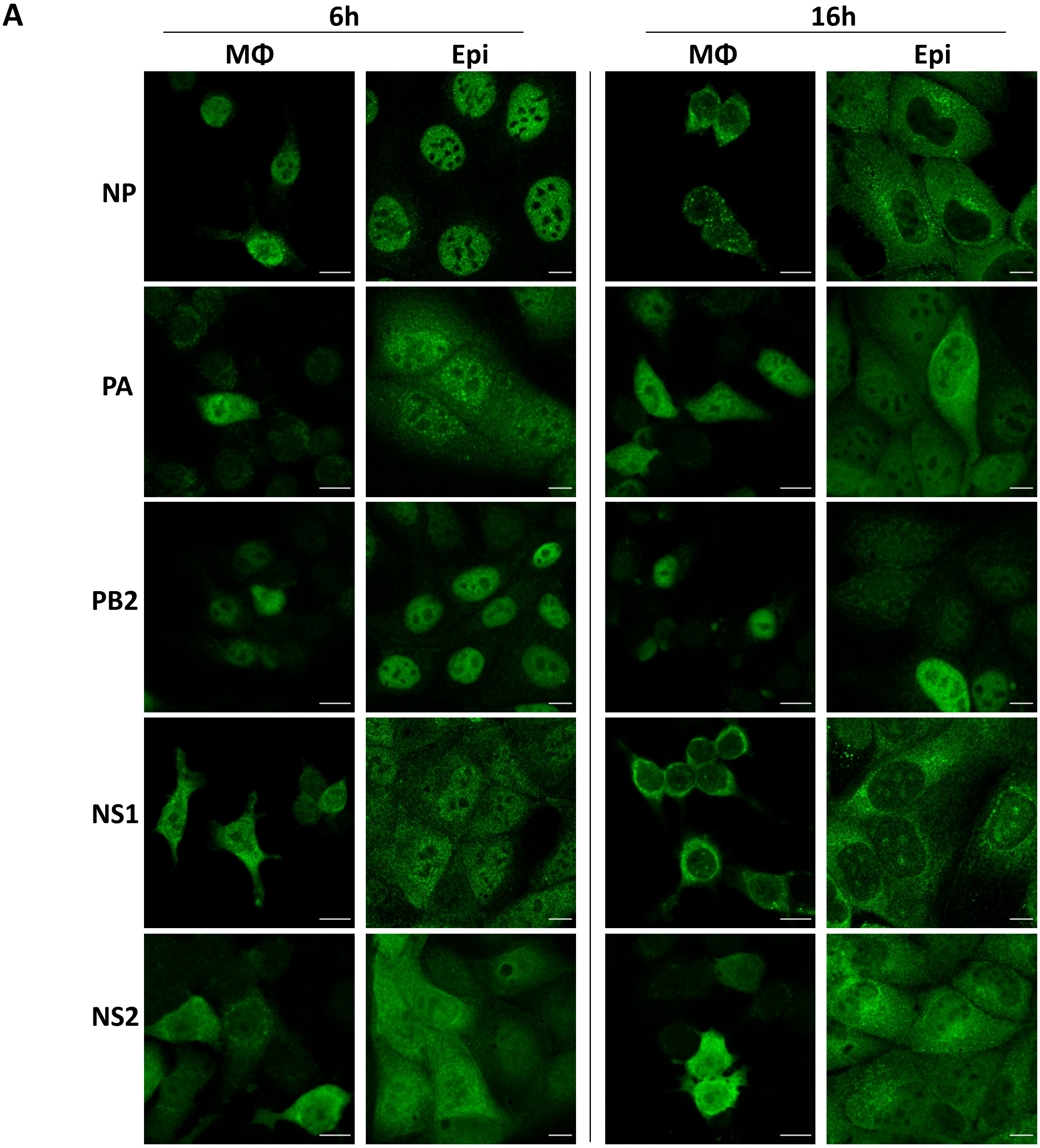


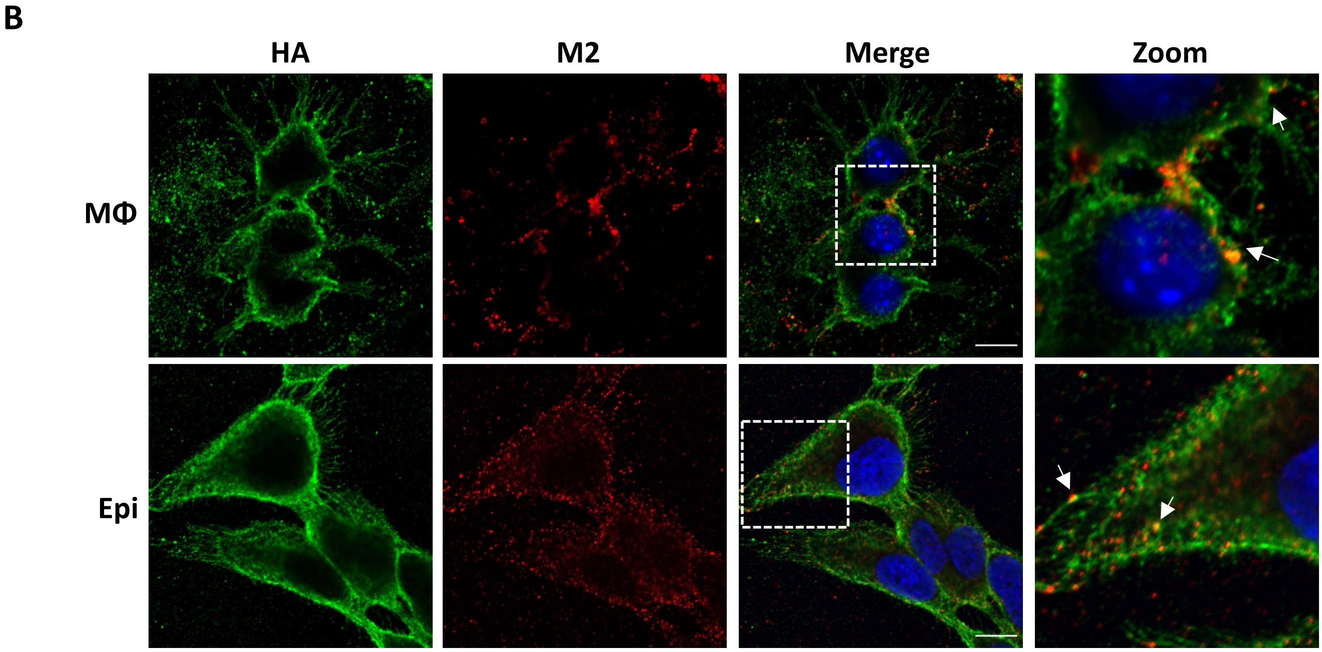

**SI 1: Viral proteins locate to similar compartments in infected MΦ compared to epithelial cells**

Mouse MΦ (RAW264.7) or epithelial cells (MDCK, as controls for productive replication) were infected with IAV (MOI 5) and fixed at 6 and 16 hpi. **(A)** Fixed cells were stained for the viral proteins NP, PA, PB2, NS1 and NS2. Scale bars = 10µM. **(B)** Mouse MΦ (RAW264.7) or epithelial cell controls (MDCK) were infected with IAV (MOI 5) and fixed at 16 hpi. Cells were stained on the cell surface for the viral proteins HA and M2. Arrows highlight areas of co-localisation in zoomed in merged image.

Suppl. Figure 2

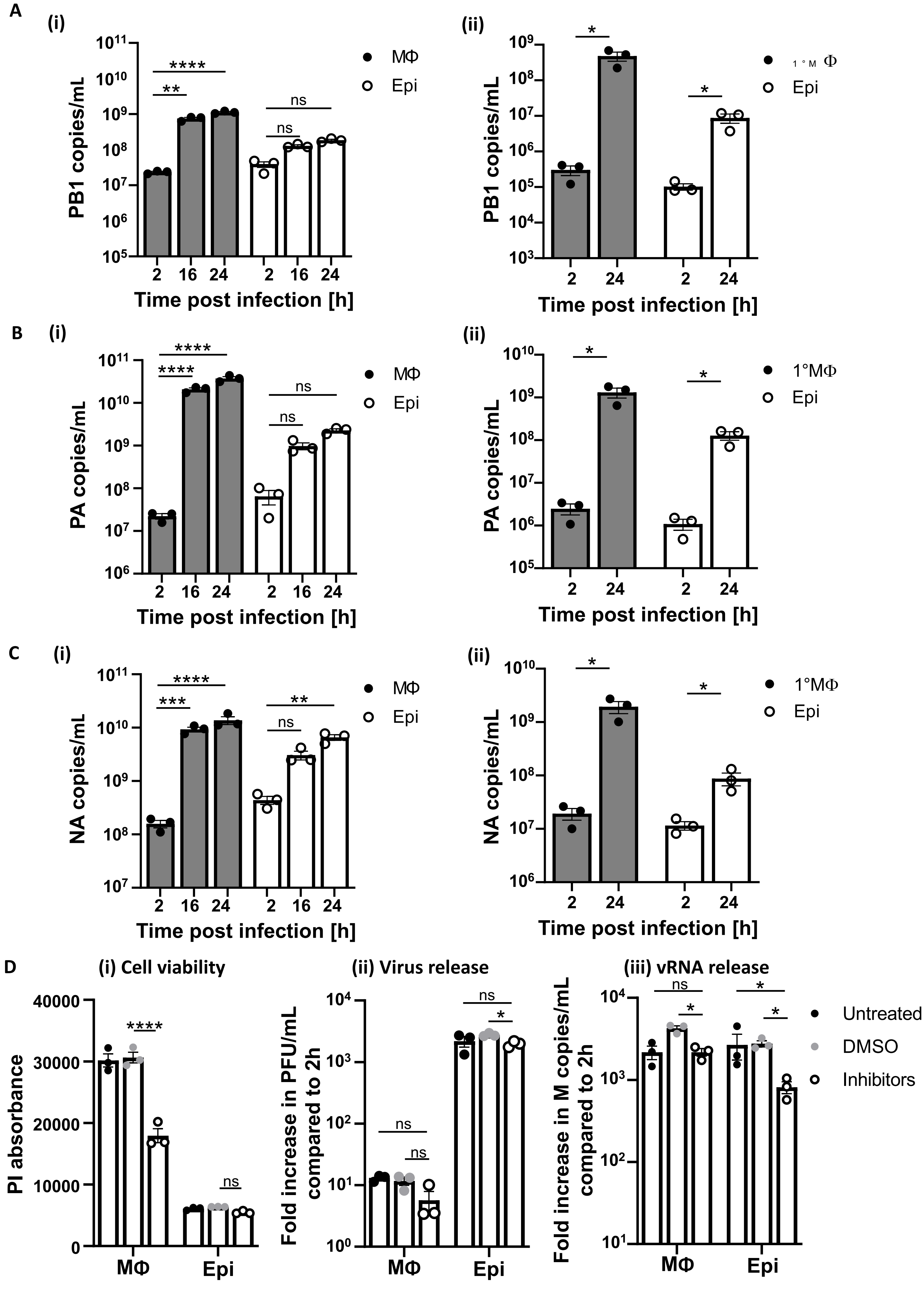

**SI 2: IAV infected MΦ release vRNAs during abortive replication**

Mouse MΦ (RAW264.7) or epithelial cell controls (MDCK) were infected with IAV (MOI 5) and SN samples were collected at 2, 16 and 24 hpi. RNA was extracted from cell culture SN and vRNA-specific cDNA was generated. The following vRNA segments were analysed via RT-qPCR to determine the copy number per mL **(A) (i)** PB1, **(B) (i)** NA, **(C) (i)** PA. Primary mouse MΦ (PEC) or epithelial control cells (LA-4) were infected with IAV (MOI 5) and the presence of the following vRNA segments was analysed via RT-qPCR at 2 or 24 hpi: **(A) (ii)** PB1, **(B) (ii)** NA, **(C) (ii)** PA. All experiments n=3; t-test or ANOVA (where applicable) was performed with $p<0.5$ *, $p<0.1$ **, $p<0.01$ ***, $p<0.001$ ****, ns not significant. **(D)** Mouse MΦ (RAW264.7) or epithelial cell controls (MDCK) were infected with IAV (MOI 5) and either left untreated or treated with DMSO control or both 10 µM Q-VD-OPh (apoptosis inhibitor) and 2 µM GSK-872 (necroptosis inhibitor) and analysed for (i) cell viability through propidium iodide (PI, 6 µg/mL) uptake at 24 hpi, (ii) IAV release at 24 hpi via plaque assay and (iii) vRNA release from infected cells at 24 hpi measured by M vRNA copies.

Suppl. Figure 3

**A**

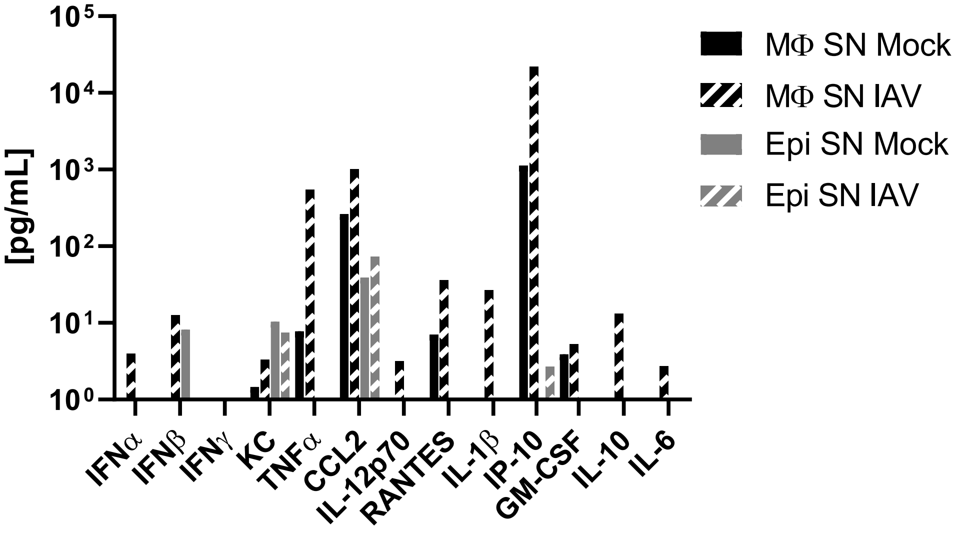


**B**

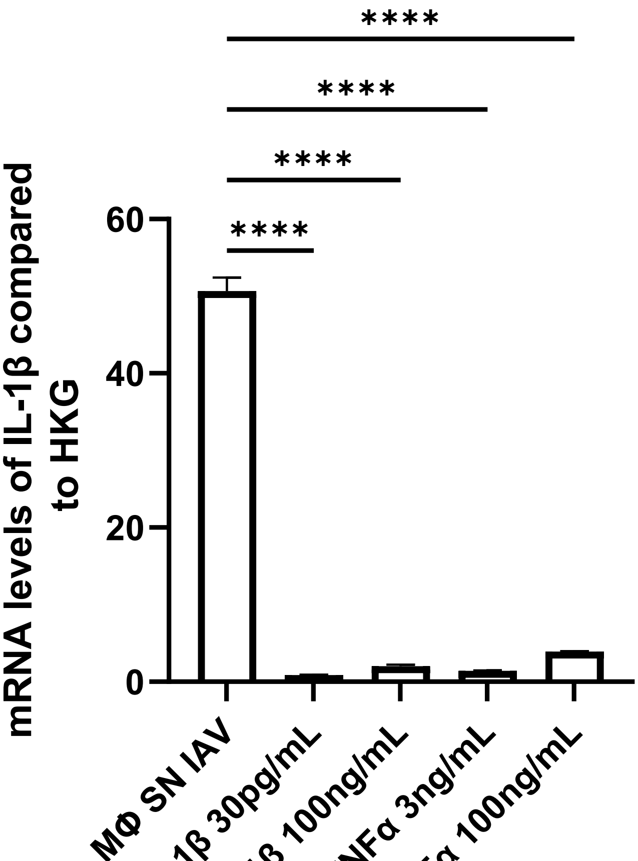

**SI 3: Selected cytokine and chemokines in the SN of infected MΦ and epithelial cells**

**(A)** Virus-cleared SN from infected and mock infected MΦ and epithelial cells was analysed for the presence of IFNα, IFNβ, IFNγ, KC, TNFα, CCL2, IL-12p70, RANTES, IL-1β, IP-10/CXCL10, GM-CSF, IL10 and IL-6 through cytometric bead array (CBA) and expressed as pg/ml. **(B)** THP-1 monocytes were either left untreated, treated with MΦ IAV SN, mouse IL-1β as detected in MΦ infected SN (30 pg/mL) or a higher concentration (100ng/mL) or mouse TNFα as detected in MΦ infected SN (3 ng/mL) or at a higher concentration (100ng/mL) for 6h and then analysed for the expression of IL-1β mRNA via RT-qPCR.
